## Supplementary figures for "Early changes in the properties of CA3 engram cells explored with a novel viral tool"

### Figure S1

A

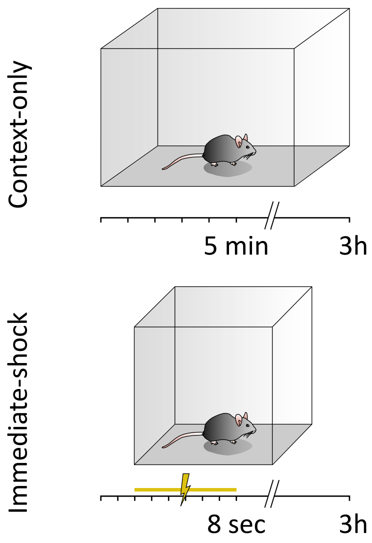
Context-only
5 min
3h
Immediate-shock
8 sec
3h

B

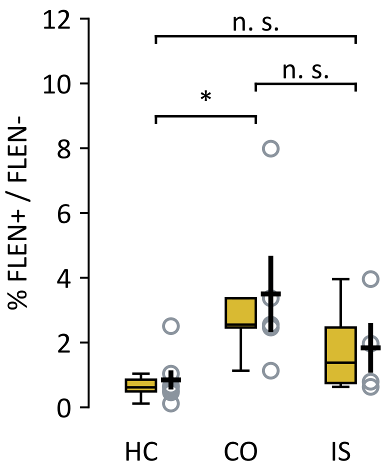
12
10
8
6
4
2
0
% FLEN+ / FLEN-
n. s.
n. s.
*
HC
CO
IS

### Figure S2

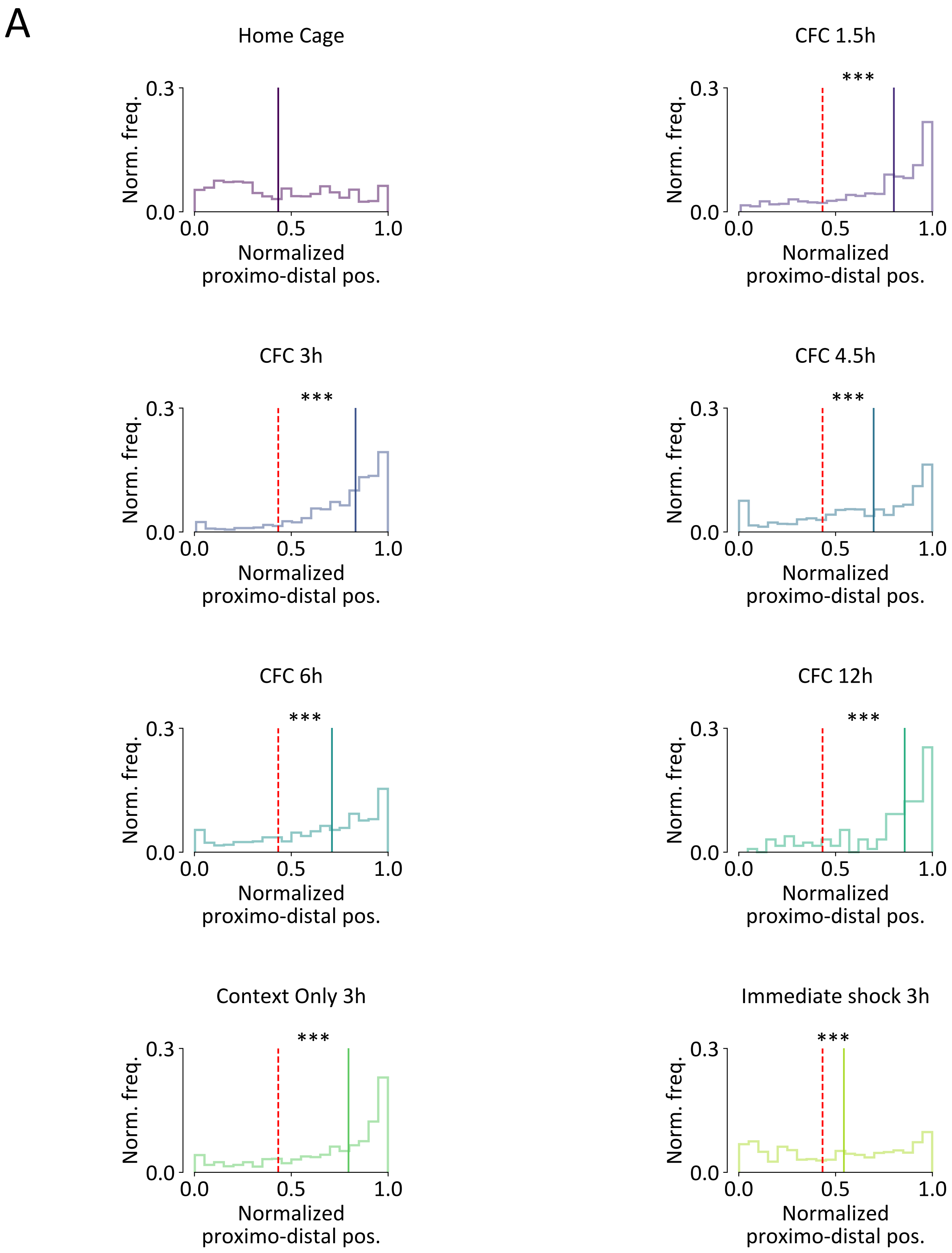
A
Home Cage
CFC 1.5h
CFC 3h
CFC 4.5h
CFC 6h
CFC 12h
Context Only 3h
Immediate shock 3h
***
Norm. freq.
0.0
0.3
0.0
0.5
1.0
Normalized
proximo-distal pos.

### Figure S3

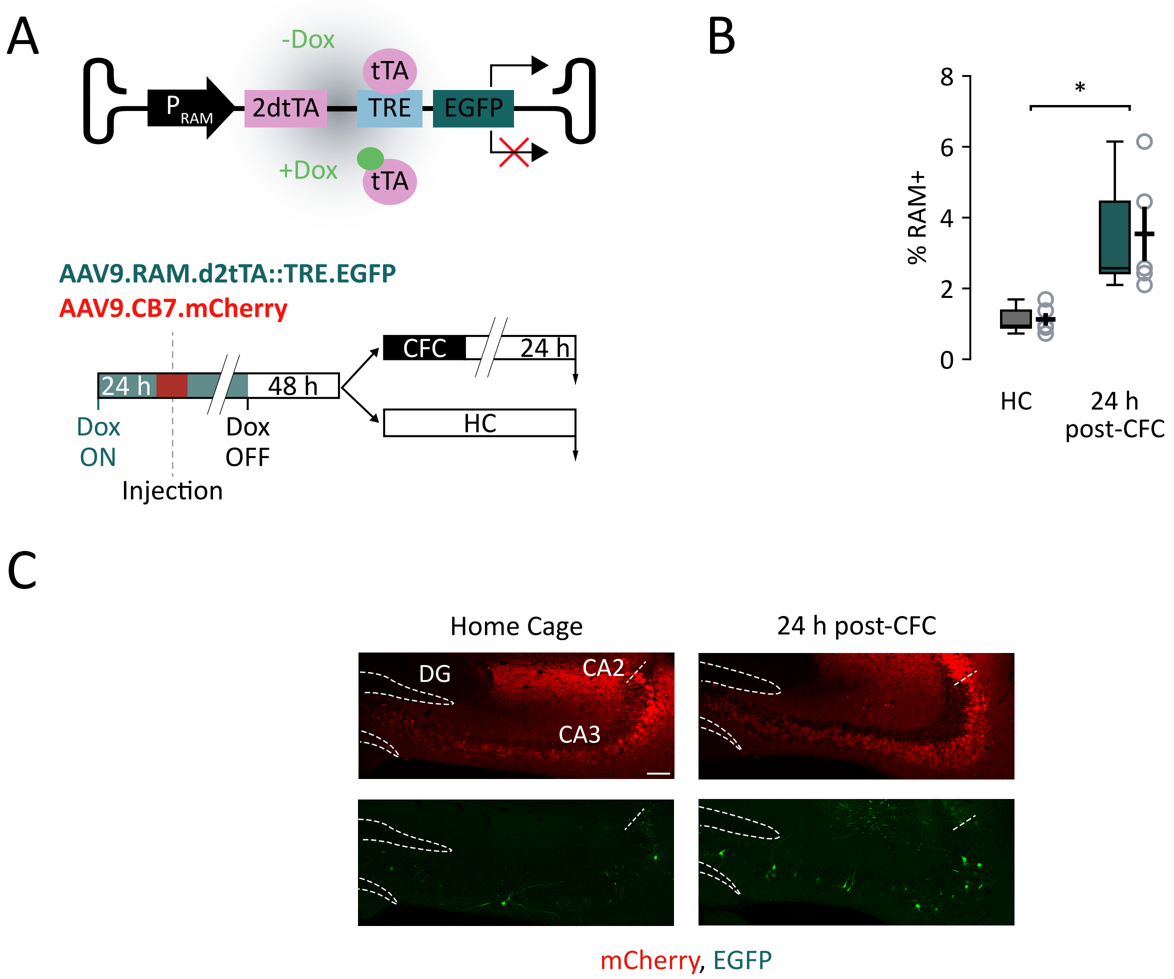
A
-Dox
tTA
P RAM
2dtTA
TRE
EGFP
+Dox
tTA
AAV9.RAM.d2tTA::TRE.EGFP
AAV9.CB7.mCherry
CFC
24 h
24 h
48 h
HC
Dox
ON
Dox
OFF
Injection
B
% RAM+
8
6
4
2
0
*
HC
24 h
post-CFC
C
Home Cage
24 h post-CFC
DG
CA2
CA3
mCherry, EGFP

### Figure S4

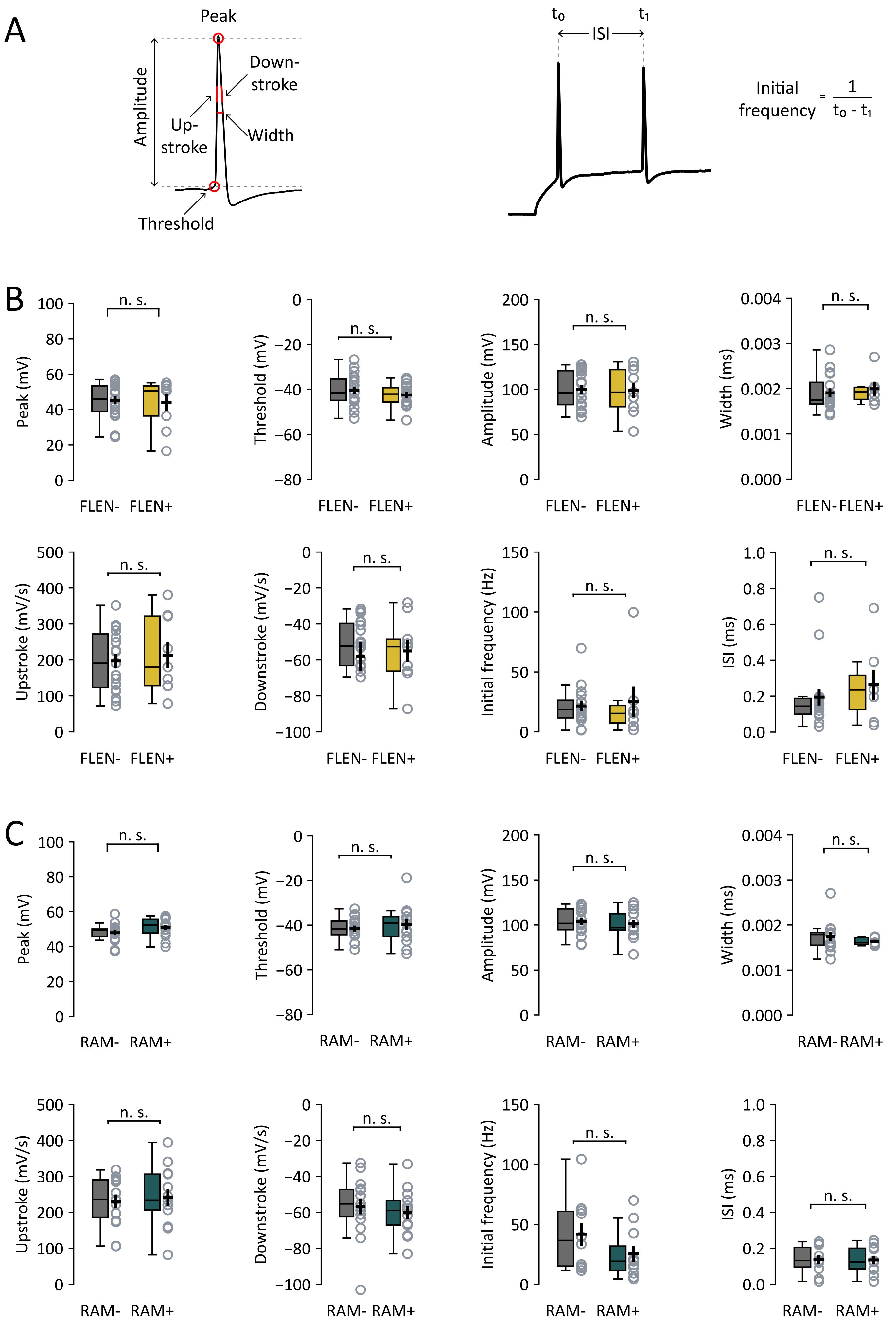
A
Peak
Amplitude
Down-
stroke
Up-
stroke
Width
Threshold
t0
t1
ISI
Initial frequency = 1 / t0 - t1
B
Peak (mV)
Threshold (mV)
Amplitude (mV)
Width (ms)
Upstroke (mV/s)
Downstroke (mV/s)
Initial frequency (Hz)
ISI (ms)
n. s.
FLEN-
FLEN+
C
RAM-
RAM+

**Figure S1.** (**A**) Left panel: context-only and immediate-shock behavioral test layout. While in the context-only task mice freely explore an arena without being conditioned, mice subjected to the immediate-shock test experience a long (8 seconds) foot-shock immediately after being placed in the conditioning cage, preventing them exploring the environment and forming contextual memories. Right panel: plot comparing the percentage of FLEN+ cells in the home cage group (from Fig. 2D) and the context-only and immediate-shock groups.

**Figure S2.** (**A**) Normalized proximodistal (x) and superficial-to-deep (y) frequency distribution of CA3 FLEN+ cells position in all behavioral groups. Black dashed lines indicate each group's median value, while red dashed line represents the home cage group's median value. Each plot includes the scattered x-y position of FLEN+ cells within the pyramidal layer, and the frequency distribution along the proximodistal (top) and superficial-to-deep (right) axis.

**Figure S3.** (**A**) RAM construct outline and expression mechanism. Top panel: Dox administration prevents binding of tTA protein to the TRE promoter, while Dox deprivation allows downstream EGFP transcription. Bottom panel: *in vivo* experiment schema for RAM system. Dox-fed mice are bilaterally injected in CA3 with RAM and infection marker AAV-CB7-mCherry. The two groups of mice (CFC vs. HC) underwent all experimental steps in parallel. Specifically, both groups were maintained on and off Dox for the same duration and received viral injection on the same day. 48 hours after Dox withdrawal, the CFC group was trained for contextual conditioning, while the HC group remained in the home cage in the holding room. All animals were thus sacrificed 72 hours after Dox removal. (**B**) Percentage of RAM+ CA3 neurons over the total number of mCherry+ neurons in HC and 24 hours post-CFC. (**C**) Representative sections of CFC-trained mice compared to HC untrained mice. Dashed lines outline the dentate gyrus (DG) cellular layer, while the dashed segment indicates the separation between CA3 and CA2. Scalebar = 100 μm.

**Figure S4.** (**A**) Left panel: overview of action potential features analyzed. Right panel: Initial frequency calculation. (**B** and **C**) Summary of analyzed action potential properties in FLEN+ vs. FLEN- (B) and RAM+ vs. RAM- (C).
